# In-frame deletion of SPECC1L microtubule binding domain results in embryonic tissue movement and fusion defects

**DOI:** 10.1101/2021.01.28.428634

**Authors:** Jeremy P. Goering, Luke W. Wenger, Marta Stetsiv, Michael Moedritzer, Everett G. Hall, Dona Greta Isai, Brittany Jack, Zaid Umar, Madison K. Rickabaugh, Andras Czirok, Irfan Saadi

## Abstract

Embryonic morphogenesis of the neural tube, palate, ventral body wall and optic fissure require precise sequence of tissue movement and fusion, which if incomplete, leads to anencephaly/exencephaly, cleft palate, omphalocele and coloboma, respectively. These are genetically heterogeneous birth defects, so there is a continued need to identify etiologic genes. Patients with autosomal dominant *SPECC1L* mutations show syndromic malformations, including hypertelorism, cleft palate and omphalocele. These *SPECC1L* mutations cluster in the second coiled-coil domain (CCD2), which facilitates association with microtubules. To study SPECC1L function in mice, we first generated a null allele (*Specc1l^ΔEx4^*) lacking the entire SPECC1L protein. Homozygous mutants for these truncations died perinatally without cleft palate or exencephaly. Given the clustering of human mutations in CCD2, we hypothesized that targeted perturbation of CCD2 may be required. Indeed, homozygotes for in-frame deletions involving CCD2 (*Specc1l^ΔCCD2^*) resulted in ~50% exencephaly and ~50% cleft palate. Interestingly, these two phenotypes are never observed in the same embryo. Examination of embryos with and without exencephaly revealed that the oral cavity was narrower in exencephalic embryos, which allowed palatal shelves to elevate despite their defect. In contrast to an evenly distributed subcellular expression pattern, mutant SPECC1L-ΔCCD2 protein showed abnormal subcellular localization, decreased overlap with microtubules, increased actin bundles, and dislocated non-muscle myosin II to the cell cortex. Thus, we show that perturbations of CCD2 in the context of full SPECC1L protein affects tissue fusion dynamics, indicating that human *SPECC1L* CCD2 mutations are gain-of-function. Improper SPECC1L subcellular localization appears to disrupt connections between actomyosin and microtubule networks, which in turn may affect cell alignment and coordinate movement during tissue morphogenesis.

## Introduction

Many embryonic tissue morphogenetic events require movement and fusion events. Examples include neural tube, palate, ventral body wall and optic fissure closure (Martin & Wood, 2002; Ray & Niswander, 2012). These processes require a precise sequence of remodeling events in a narrow and time-sensitive window of embryogenesis. Incorrect movement or incomplete fusion in the three examples described above leads to structural birth defects of anencephaly/exencephaly (1/4000 live births), cleft palate (1/1700), omphalocele (1/4000) and coloboma (1/2000) respectively (Mai et al., 2019).

We are studying the cytoskeletal protein SPECC1L that is proposed to associate with both actin filaments and microtubules via its calponin homology domain (CHD) and coiled coil domain 2 (CCD2), respectively (Hall et al., 2020; Saadi et al., 2011; Wilson et al., 2016). We and others have shown that patients with autosomal dominant *SPECC1L* mutations manifest a range of structural birth defects, including hypertelorism (97%), omphalocele (45%) and cleft palate (24%) (Bhoj et al., 2018; Bhoj et al., 2015; Kruszka et al., 2015; Saadi et al., 2011). Interestingly, all *SPECC1L* mutations in these patients with syndromic manifestation cluster largely in CCD2 and to a lesser extent in CHD, indicating a critical role for these domains.

To understand the function of SPECC1L in the etiology of these structural defects, we generated and evaluated several different mouse alleles, including a C-terminal truncation that also removed the CHD (*Specc1l^ΔC510^*) (Hall et al., 2020) and null allele lacking any protein (*Specc1l^ΔEx4^,* Figure 1). Homozygous mutants for these alleles exhibited perinatal lethality but did not show cleft palate or omphalocele (Hall et al., 2020). We did observe a delay in palatal shelf elevation in *Specc1l^ΔC510^* compound heterozygotes accompanied by both palatal shelf epithelial (Hall et al., 2020) and mesenchymal defects (Goering et al., 2021). Thus, we hypothesized that patient phenotypes of cleft palate and omphalocele were due to gain-of-function CCD2 mutations.

**Figure 1.**
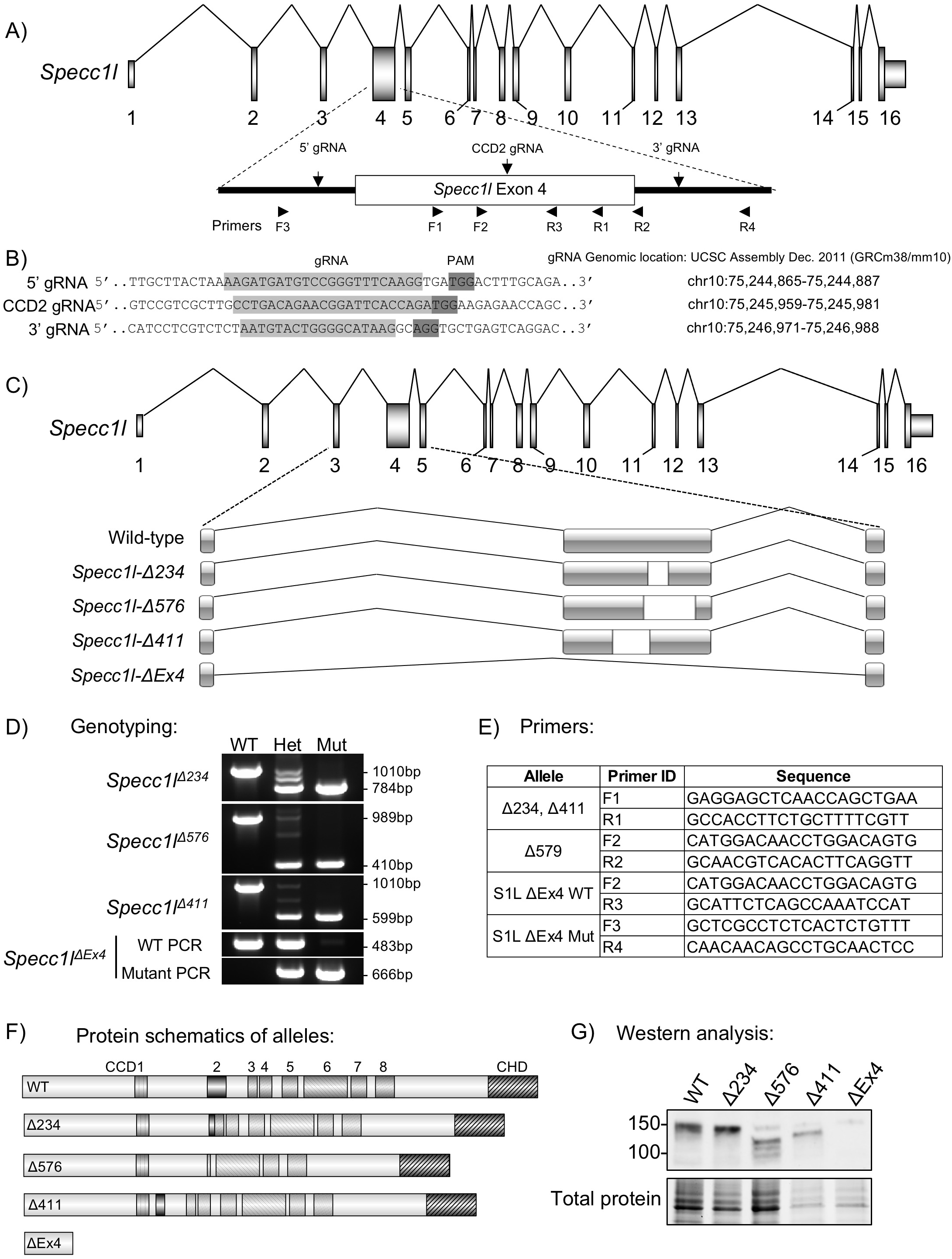
Generation of *Specc1l* null and CCD2 deletion alleles. **A)** Schematic of *Specc1l* locus. The largest exon 4, which also encodes the coiled-coil domain 2 (CCD2), is highlighted. The locations of the two guide RNAs (5’ and 3’ gRNAs) used to delete exon 4, as well as the gRNA used to introduce deletions in CCD2 (CCD2 gRNA) are indicated. Also shown are the approximate locations of the sequencing primers used for genotyping in panels D and E. **B)** Sequences and genomic locations of the gRNAs used in panel A. PAM = Protospacer Adjacent Motif. **C)** Schematic representation of the genomic deletions created by the three CCD2 in-frame (Δ234, Δ576, Δ411 base pairs) and the null exon 4 deletion (ΔEx4). **D)** Genotyping analysis of the four alleles, showing heterozygous and homozygous occurrence of the slower migrating band corresponding to deletion. Due to the large deletion in ΔEx4, wildtype and mutant alleles are genotyped separately. **E)** Sequence for primer pairs used for genotyping shown in panel D. Primer locations are also shown in the schematic of exon 4 in panel A. **F)** SPECC1L protein schematics resulting from the CCD2 deletions and the null allele. CCD = coiled-coil domain; CHD = calponin homology domain. **G)** Western blot analysis of tissue from wildtype (WT) and homozygous mutants for the four alleles. WT SPECC1L band migrates at ~150 KDa. For the three in-frame CCD2 deletion alleles (Δ234, Δ576, Δ411), corresponding faster migrating bands are visible. There are no bands visible for the ΔEx4 null allele. Total protein is shown for loading control.

To test this hypothesis, we targeted CCD2 with CRISPR based genome editing. Here we report three alleles with in-frame genomic deletions removing portions of CCD2 (collectively termed *Specc1l^ΔCCD2^).* Homozygous mutants for all three alleles showed either cleft palate or exencephaly, and omphalocele at high penetrance. In contrast, homozygous mutants for *Specc1l^ΔEx4^* null allele showed none at birth. In addition, we observed significantly higher penetrance of cleft palate in female *Specc1l^ΔCCD2/+^* heterozygotes than male counterparts. We also provide cellular and molecular evidence for a role of SPECC1L in regulating actomyosin organization during tissue morphogenesis.

## Materials and Methods

### Generation of Specc1l alleles

The general scheme for the CRISPR based generation of *Specc1l* alleles is described in **Figure 1**. Briefly, to generate the *ΔEx4* null allele, two gRNAs flanking exon 4 (**Figure 1A,B**) were introduced directly in F1 hybrid (C57Bl/6J:FVB/NJ) zygotes at 2-4 cell stage. Successful targeting was confirmed in the founders using PCR based loss of exon 4. The CCD2 specific alleles were generated using a single gRNA located in CCD2 in exon 4 (**Figure 1A,B**). Progeny from founders were sequenced for in-frame deletions. Three of the in-frame deletions (Δ234, Δ576, Δ411 base pairs) were established as mouse lines and further analyzed (**Figure 1C-G**).

### Mouse embryo processing and histological analysis

Timed-matings were set up overnight and checked for plugs the following morning. The embryos were classified as embryonic day 0.5 (E0.5) at noon on the day that the plug was identified. At the desired embryonic time point, the pregnant females were euthanized using methods approved by IACUC. Embryos were harvested, washed in 1x PBS, and fixed overnight in 4% PFA. Prior to fixation, whole embryo brightfield images were obtained using a Nikon SMZ 1500 stereomicroscope. Yolk sacs were taken at the time of harvesting for genotyping. Sex was determined by PCR using primers flanking an 84 bp deletion of the *Rbm31x* gene relative to its gametolog *Rbm31y* (Forward: CACCTTAAGAACAAGCCAATACA; Reverse: GGCTTGTCCTGAAAACATTTGG). A single product (269 bp) was amplified in female subjects and two products (269 bp; 353 bp) amplified in male subjects (Tunster, 2017). Whole mount DAPI staining of the palate was achieved by decapitating fixed embryos, removing the lower jaw, and incubating the exposed palates in 500-1000nM DAPI solution overnight, then imaging on Nikon SMZ 1500 stereomicroscope. Fixed embryo heads were paraffin processed using Leica ASP300, embedded in paraffin blocks, and coronally sectioned. H&E staining was performed on these sections and imaged using the EVOS FL Auto microscope. For cryosectioning, fixed embryo heads were submerged in 20% sucrose until the tissue sank, embedded in OCT media, coronally sectioned at a thickness of 10 μm, then stored at −80°C until immunostained.

### Mouse embryonic palatal mesenchyme (MEPM) isolation and culture

MEPM isolation was performed as described previously (Goering et al., 2021). Briefly, harvested E13.5 mouse embryos were decapitated, the lower jaw and tongue were removed, the palatal shelves were excised, and incubated in 0.25% trypsin (ThermoFisher, 25200056) at 37°C for 10 minutes. The resulting MEPM cells were resuspended in DMEM high glucose supplemented with 10% Fetal Bovine Serum (Corning, 35-010-CV), 40mM L-Glutamine (Cytiva, SH30243.01) and 50 Units/mL penicillin/streptomycin (Corning, 30-002-CI). MEPMs were plated into 6-well plates and incubated at 37°C with 5% CO2. Culture media was changed daily until the cells reached confluency, whereupon they were cryopreserved. For immunostaining, cells were thawed and seeded at the appropriate density.

### Spontaneous collective motility assay

For the analysis of stream formation and collective motility, MEPM cells were seeded at high (450-900/mm^2^) densities into selected areas of 6 well plates, delimited by silicone insert rings (Ibidi, 80209). Cultures were live imaged every 10 minutes with a 4x phase contrast objective. The local spatial correlations of cell movements were characterized by the average flow field that surrounds moving cells as described in (Czirok, Varga, Mehes, & Szabo, 2013; Goering et al., 2021; Szabo et al., 2010). Briefly, cell motility was estimated using a particle image velocimetry (PIV) algorithm, with an initial window size of 50 μm (Czirok et al., 2017; Goering et al., 2021; Zamir, Czirok, Rongish, & Little, 2005), resulting in velocity vectors at each frame and image location. A reference system was then aligned to each vector and adjacent vectors were registered in the appropriate bin (front, rear, etc.). The average velocity vector in each bin was used as a measure of spatial correlation. The calculated average co-movement vectors were fitted with an exponential function across the front-rear and left-right axes to obtain the correlation length: the characteristic distance local correlations in cell velocity disappear.

### Western blotting

Protein was extracted by sonicating embryonic tissue in radioimmunoprecipitation buffer with HALT protease inhibitor cocktail (Thermo Scientific, 78440). Protein concentrations were quantified using BCA assay (Thermo Scientific, 23227). Samples were electrophoresed using Mini-Protean TGX Stain-Free 4-15% gradient pre-cast polyacrylamide gels (Bio-RAD, #4568084) and transferred to an Immobilon PVDF membrane (EMD Millipore, IPVH00010). Membranes were blocked using Odyssey Blocking Buffer (Li-Cor, 927-5000) for 1-2 hours at room temperature, incubated with primary antibody overnight at 4°C, secondary antibody (1:10000; Cell Signaling Technologies) for 1 hour at room temperature, and developed using Femto Super Signal West ECL reagent (Thermo Scientific, 34095). Membranes were imaged and analyzed using ChemiDoc MP imaging system (BioRad) and Imagelab software (BioRad). Primary antibodies used: SPECC1L (1:2000; Proteintech, 25390-1-AP), MYH10 (1:1000; Sigma, M7939).

### Immunofluorescence

For MEPM immunostaining, cells were grown on poly-lysine coated glass coverslips and fixed with 2% paraformaldehyde for 10 min at RT. Cells were permeabilized with 0.1% Trixon X-100 in 1× PBS for 10 min, washed with PBS 3x, and blocked in 10% normal goat serum (NGS) (Thermo Fisher Scientific, 50062Z) for 2 hrs at room temperature. Primary antibodies were incubated overnight at 4°C, washed in 1× PBS, then incubated in secondary antibody for 2 hours and DAPI and/or F-actin stain for 30 min at room temperature. Stained coverslips were mounted on slides with Prolong Gold Antifade Mounting medium (Thermo Fisher Scientific, P10144). For ΔNp63 and β-catenin, an antigen retrieval protocol was followed by heating slides in sodium citrate buffer (10 mm Sodium Citrate, 0.05% Tween 20, pH 6.0) at 96°C for 10 min. Slides were washed in H2O and PBS, permeabilized using 0.5% Triton X-100 in 1× PBS for 30 min, washed in PBS again, then blocked in 10% NGS. Primary and secondary antibodies were incubated following the same method as described above. Images were acquired using either an EVOS FL Auto Inverted Imaging System or Leica SP8 Stimulated emission depletion (STED) 3x WLL confocal laser microscope. Primary antibodies: SPECC1L N-terminus (1:500 cells, 1:250 tissue Proteintech, 25 390-1-AP), α-tubulin (1:100, 1:500 cells, 1:1000 tissue, Sigma, T9026), ΔNp63 (1:100, Biolegend, 619001), β-catenin (1:500 Cell Signaling Technology; 2677), MYH9 NM-IIA (1:100 Proteintech, 11128-1-AP), MYH10 NM-IIB (1:100 Sigma, M7939). Secondary antibodies and stains: Goat anti-Rabbit IgG (H+L) Alexa 488 and 594 (1:1000 cells, 1:500 tissue Invitrogen; A-11008, A-11012), Goat anti-Mouse IgG1 Alexa 488 (1:500 Invitrogen, A-21121), Goat anti-Mouse IgG1 Alexa 680 (Jackson Immuno, 1156255205), Acti-stain 555 phalloidin (1:140, Cytoskeleton, PHDH1-A), Actin-stain 670 phalloidin (1:140, Cytoskeleton, PHDN1-A), DAPI (5 μM).

### Immunoprecipitation

Protein was extracted by sonicating embryonic tissue in a lysis buffer containing 20mM Tris-HCl, 1% NP-40, 131mM NaCl, 2mM EDTA, and 10% glycerol. To prevent nonspecific binding to the protein beads, the lysate was pre-cleared by incubating with Dynabeads Protein G (Invitrogen, 100-03D) for 4 hours at 4°C while rocking. For each sample, equal volume of lysate was incubated with 1μg of either SPECC1L N-terminus antibody (Proteintech, 25390-1-AP) or Normal Rabbit IgG control antibody (Cell Signaling Technology, 2729S). Dynabeads Protein G were added to the lysate/antibody mixture and incubated overnight at 4°C while rocking. The beads were washed with ice cold lysis buffer three times. Lysis buffer containing 5X Laemmli buffer with 10% 2-Mercaptoethanol (Sigma, M3148) was added to the beads, and then electrophoresed on polyacrylamide gels as described above for Western Blotting.

## Results

### *Specc1l* null mutants show perinatal lethality with no fusion defects

We previously generated three different mouse alleles for *Specc1l* deficiency using a genetrap-based knockdown strategy (Hall et al., 2020; Wilson et al., 2016). Homozygous mutants for all three genetrap alleles showed early embryonic lethality with a neural tube closure defect (Hall et al., 2020; Wilson et al., 2016). We also generated an allele (*Specc1l^ΔC510^*) with a C-terminal truncation that deleted 510 amino acids including the CHD (Hall et al., 2020). Homozygous mutants for this allele were perinatal lethal but did not show exencephaly or cleft palate, however, the truncated protein was still expressed (Hall et al., 2020). To determine the true *Specc1l* null phenotype, we have now used CRISPR guide RNAs flanking exon 4 (**Figure 1A,B**) to generate a knockout allele lacking exon 4 (*Specc1l^ΔEx4^,* **Figure 1C**) that does not show any SPECC1L protein product (**Figure 1F,G**). Homozygous mutants for *Specc1l^ΔEx4^* are also perinatal lethal and do not show exencephaly or cleft palate (**Figure 2**). Thus, we hypothesized that CCD2 mutations resulting in cleft palate and omphalocele were gain-of-function.

**Figure 2.**
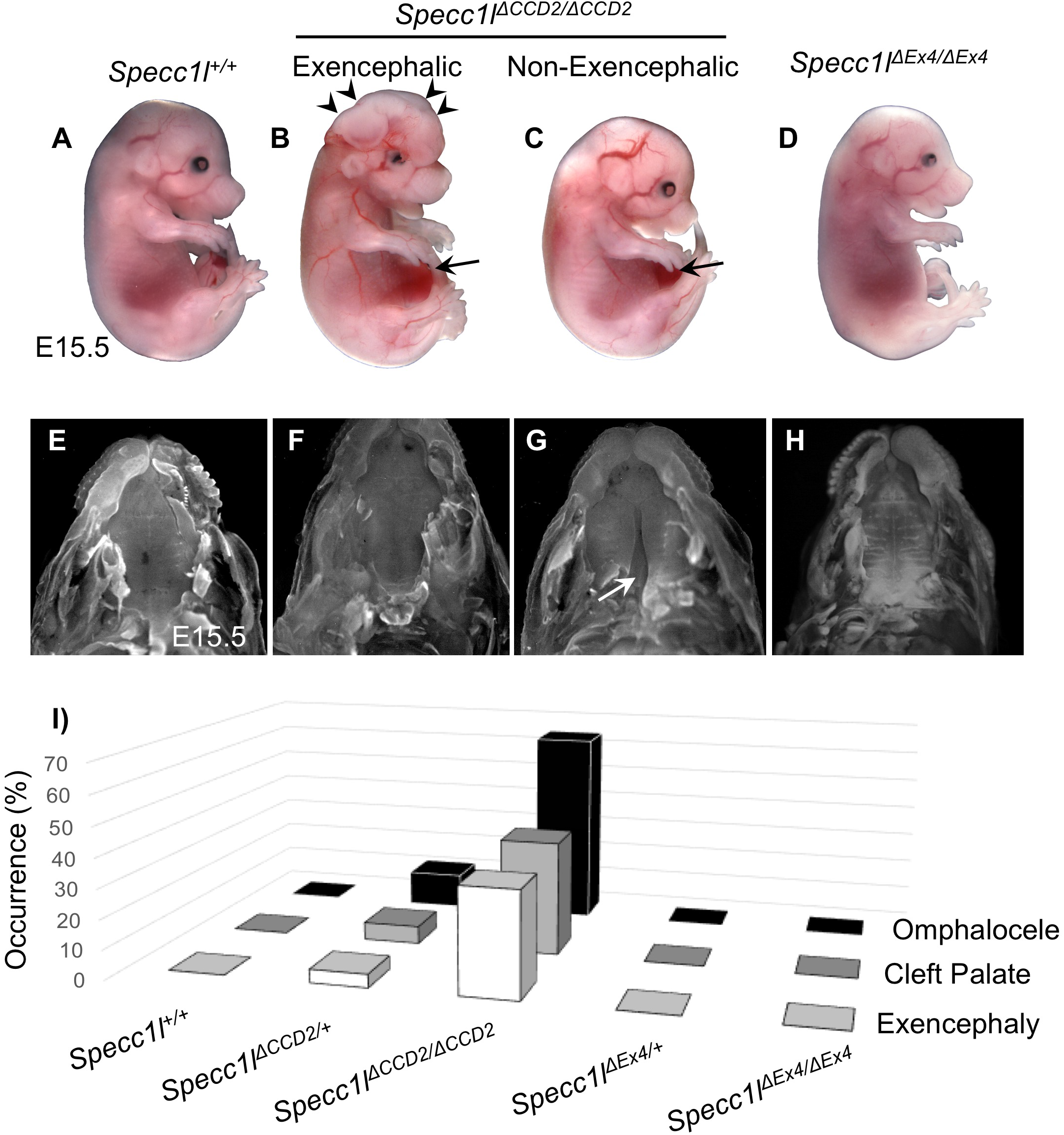
Homozygous mutants with in-frame CCD2 deletions show omphalocele, cleft palate and exencephaly. **A-D)** Gross images of wildtype (A, *Specc1l^+/+^), Specc1l^ΔCCD2/ΔCCD2^,* with (B) or without (C) exencephaly, and *Specc1l^ΔEx4/ΔEx4^* null embryos at embryonic day (E) 15.5. Exencephaly is indicated in panel B with arrowheads. Omphalocele is indicated in panels B and C with arrows. **E-H)** The palates are shown for E15.5 embryos with the jaws removed for the genotypes in panels A-D. The *Specc1l^ΔCCD2/ΔCCD2^* mutant with exencephaly (F) and the *Specc1l^ΔEx4/ΔEx4^* null (H) embryos do not show cleft palate, while the *Specc1l^ΔCCD2/ΔCCD2^* mutant without exencephaly does (G, white arrow). **I)** The prevalence (%) of omphalocele, cleft palate and exencephaly is shown in *Specc1l^ΔCCD2^* and *Specc1l^ΔEx4^* heterozygous and homozygous mutants. For this analysis, data from all three in-frame CCD2 deletion alleles (Δ234, Δ576, Δ411) were combined as ΔCCD2. The *Specc1l^+/+^, Specc1l^ΔEx4/+^,* and *Specc1l^ΔEx4/ΔEx4^* embryos did not show any occurrence of omphalocele, cleft palate or exencephaly.

### *Specc1l* mutants with in-frame deletion of CCD2 result in exencephaly, cleft palate and ventral body wall closure defects

To test this hypothesis, we targeted CCD2 with a CRISPR guide RNA (CCD2 gRNA; **Figure 1A,B**) and identified alleles with both out-of-frame and in-frame deletions. Three alleles with in-frame deletions (*Specc1l^Δ234^, Specc1l^Δ576^, Specc1l^Δ411^*) were further characterized. Genomic changes were confirmed (**Figure 1C**), and genotyping protocols were created (**Figure 1D,E**). Predicted protein level changes in ΔCCD2 proteins (**Figure 1F**) were validated for expression with western blot analysis using an N-terminal a-SPECC1L antibody (**Figure 1G**). Homozygous mutants for all alleles with out-of-frame deletions were perinatal lethal and showed no exencephaly, cleft palate or omphalocele at birth (data not shown), similarly to homozygous mutants for *Specc1l^ΔC510^* (Hall et al., 2020) and *Specc1l^ΔEx4^* (**Figure 2**). However, homozygous mutants for all alleles with an in-frame deletion in CCD2 - *Specc1l^Δ234^, Specc1l^Δ576^, Specc1l^Δ411^* are collectively referred to as *Specc1l^ΔCCD2^* - showed ~35% anterior neural tube closure defect (exencephaly, **Figure 2B, arrowheads**), ~38% cleft palate (**Figure 2G**), and ~65% ventral body wall closure defect (omphalocele; **Figure2B,C, arrows**). Embryonic penetrance values for all *Specc1l^ΔCCD2^* alleles are graphed collectively in **Figure 2I**, and the values for individual alleles are provided in **Supplemental Table 1**. Closer inspection of the embryos at E15.5 also showed coloboma or incomplete fusion of the optic fissure in *Specc1l^ΔCCD2^* mutants (**Figure S1B,C** vs. **S1A,D**; arrows). Thus, in-frame perturbation of CCD2 in the context of full-length SPECC1L was more detrimental to embryonic tissue movement and fusion events than complete loss of SPECC1L protein, indicating a gain-of-function mechanism.

### Homozygous *Specc1l^ΔCCD2^* mutant embryos did not simultaneously show exencephaly and cleft palate

In our phenotypic analysis, we noted that exencephaly (n=15) and cleft palate (n=17) never simultaneously occurred in the same *Specc1l^ΔCCD2/ΔCCD2^* embryo. In contrast, ventral body wall closure defect accompanied both exencephaly (n=10/15, 67%) and cleft palate (n=5/17, 29%). Histologic analysis of embryos showed that at E13.5, *Specc1l^ΔCCD2/ΔCCD2^* mutants with exencephaly had a narrower oral cavity (**Figure 3B**) than either wildtype control (**Figure 3A**), homozygous *Specc1l^ΔCCD2/ΔCCD2^* mutants without exencephaly (**Figure 3C**) or homozygous null *Specc1l^ΔEx4/ΔEx4^* mutants (**Figure 3D**). In addition to the narrow oral cavity, the palatal shelves themselves were abnormally wide, deforming the shape of the tongue (**Figure 3B, E13.5, arrowheads**). At E15.5, *Specc1l^ΔCCD2/ΔCCD2^* mutants with exencephaly are able to complete palatal shelf elevation and fusion (**Figure 3B**), albeit the oral cavity remains narrower (**Figure 3B, E15.5, arrowheads**) than those in wildtype (**Figure 3A**) or *Specc1l^ΔEx4/ΔEx4^* mutants (**Figure 3D**). Most homozygous *Specc1l^ΔCCD2/ΔCCD2^* mutants without exencephaly showed permanent cleft palate (**Figure 3C, arrow**). *Specc1l^ΔCCD2/ΔCCD2^* mutants did not survive postnatally even those without exencephaly or cleft palate. All mutants showed subtle abnormalities in palatal shelf rugae formation at E18.5 (**Figure 3C, E18.5**). Interestingly, *Specc1l^ΔEx4/ΔEx4^* mutants showed the most drastic effect on palatal rugae formation, indicating an overall defect in palatogenesis (**Figure 3D, E18.5**).

**Figure 3.**
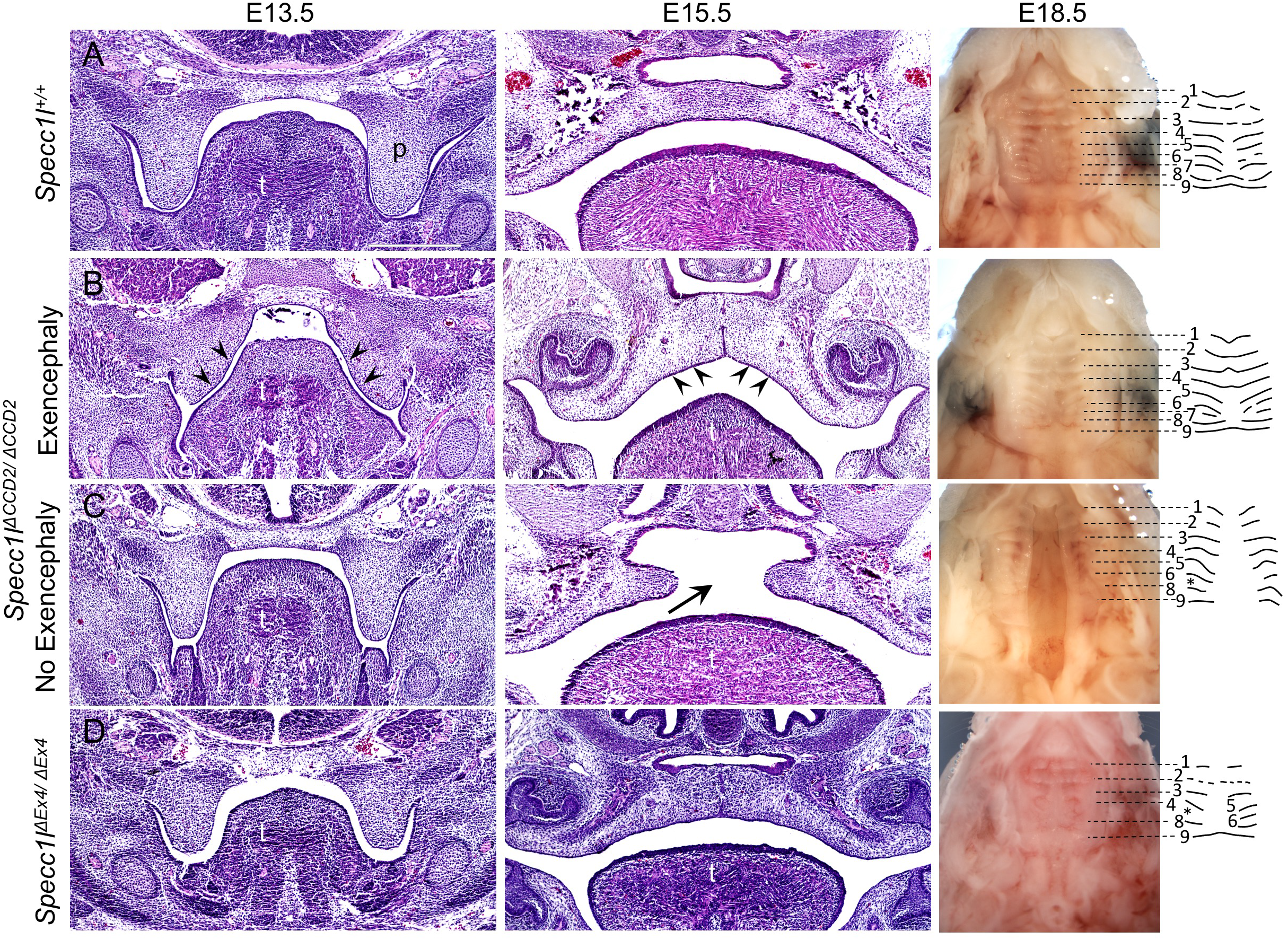
Exencephaly narrows the oral cavity allowing *Specc1l^ΔCCD2/ΔCCD2^* mutant palatal shelves to fuse. Wildtype (A), *Specc1l^ΔCCD2/ΔCCD2^* mutant with exencephaly (B), *Specc1l^ΔCCD2/ΔCCD2^* mutant without exencephaly (C) and *Specc1l^ΔEx4/ΔEx4^* null (D) embryos at embryonic day (E) 13.5, 15.5, and 18.5 are shown. At E13.5 and E15.5, coronal sections through the oral cavity were stained with hematoxylin and eosin. In the *Specc1l^ΔCCD2/ΔCCD2^* mutant with exencephaly (B) at E13.5, it is clear to see a narrowing of the oral cavity and the tongue (B, E13.5, arrowheads). Even at E15.5, the exencephalic mutant shows a narrowed dome-shaped palate and tongue (B, E15.5, arrowheads). At E18.5, embryonic palates are shown with the lower jaw removed. In addition, palatal rugae pattern at E18.5 is shown schematically. p = palatal shelf; t = tongue; * = missing palatal rugae.

### Heterozygotes for *Specc1l^ΔCCD2^* show autosomal dominant phenotypes and sex differences

The *Specc1l^ΔCCD2^* alleles were generated on a mixed C57BL/6J and FVB/NJ background. As we backcrossed the alleles to the C57BL/6J background, we noticed an occurrence of ventral body wall closure defect (12%), cleft palate (6%) and exencephaly (4.5%) in the heterozygotes (**Figure 2I**, **Table S1**). This is consistent with the autosomal dominant manifestation of human *SPECC1L* CCD2 mutations. Interestingly, so far we have only identified cleft palate in female *Specc1l^ΔCCD2/+^* heterozygotes (16%) and not in male heterozygotes (0%) (**Figure S2**). This sex difference among heterozygotes was borderline for ventral body wall closure defects (0% female vs. 5% male), and not observed for exencephaly (5% vs. 5%) (**Figure S2**). Interestingly, as we further backcrossed the alleles onto C57BL/6J to ~N6, we noticed that we were rarely getting any female heterozygotes at weaning (data not shown). We are therefore maintaining the colony on a mixed C57BL/6J and FVB/NJ background. We also looked at male to female ratios in homozygous *Specc1l^ΔCCD2^* mutants and found a small prevalence of ventral body wall closure defects in female mutants (77% female vs. 50% male; **Figure S2**). Again, a difference was not observed in homozygous *Specc1l^ΔCCD2^* mutants for cleft palate (33% female vs. 33% male) or exencephaly (55% female vs. 50% male). These results suggest an interplay between strain and sex differences along with autosomal dominance.

### Abnormal cellular expression of SPECC1L-ΔCCD2 protein in palatal shelf mesenchyme

SPECC1L protein was broadly expressed in both the palatal shelf epithelium and mesenchyme. We have previously shown that SPECC1L expression was more prominent at cell-cell boundaries in epithelial cells (Hall et al., 2020; Wilson et al., 2016). In *Specc1l^ΔCCD2/ΔCCD2^* mutant palatal shelf epithelium and periderm, we did not observe a striking difference in staining pattern or localization of SPECC1L, F-actin, or microtubules (**Figure S3**). However, in the palatal mesenchyme, we observed that SPECC1L-ΔCCD2 protein was not diffusely expressed throughout the cell and was instead concentrated towards the periphery of cells (**Figure 4A,A’** vs. **4F,F’**). We also previously showed that SPECC1L expression closely overlaps with filamentous actin (F-actin) staining (Hall et al., 2020; Wilson et al., 2016). Consistently, both wildtype and SPECC1L-ΔCCD2 proteins showed a similar co-localization with F-actin (**Figure 4E,E’** vs. **4J,J’**). However, there are many more, thick F-actin bundles in *Specc1l^ΔCCD2/ΔCCD2^* mutant palatal mesenchyme (**Figure 4F’,J’; arrows**). In contrast, SPECC1L-ΔCCD2 protein shows less colocalization with microtubules (**Figure 4D,D’** vs. **4I,I’**). These results are consistent with our previous assertion that SPECC1L CCD2 helps the protein associate with microtubules (Hall et al., 2020; Kruszka et al., 2015; Saadi et al., 2011). Thus, we argued that the gain-of-function underlying the palate elevation defect in *Specc1l^ΔCCD2/ΔCCD2^* mutants mainly affected palatal shelf mesenchyme.

**Figure 4.**
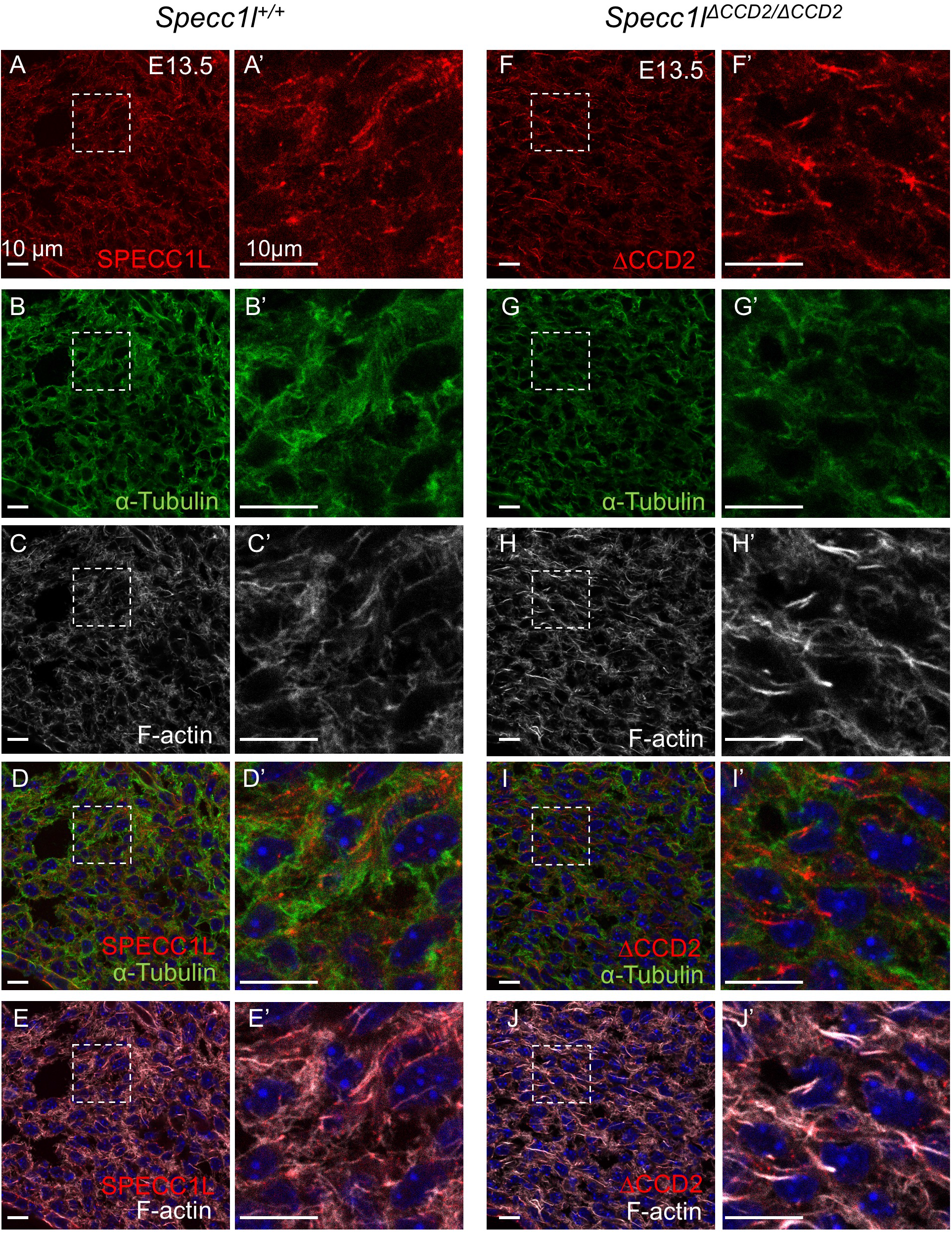
Abnormal cellular expression of SPECC1L-ΔCCD2 in E13.5 palatal shelf mesenchyme. Wildtype *Specc1l^+/+^* (A-E, A’-E’) and *Specc1l^ΔCCD2/ΔCCD2^* mutant (F-J, F’-J’) embryonic day (E) 13.5 cryosections were co-stained with antibodies against SPECC1L (A, A’, F, F’) and a-Tubulin (B, B’, G, G’), and with phalloidin (C, C’, H, H’). Panels A’-J’ are magnified regions that are boxed in the corresponding panels A-J. Wildtype SPECC1L (A, A’) shows a diffused subcellular expression that strongly overlaps with both microtubules (D, D’) and filamentous actin (E, E’). In contrast, mutant SPECC1L-ΔCCD2 (F, F’) shows an abnormal expression pattern that appears to be clustered towards cell periphery (arrows). This clustered expression of the mutant SPECC1L shows reduced colocalization with microtubules (I, I’, increased red staining), and results in abnormal bundles of F-actin (J, J’). Scalebars = 10 μm.

### CCD2 is required for subcellular diffusion of SPECC1L

To further assess the subcellular mesenchymal effects of SPECC1L-ΔCCD2 protein, we isolated primary mouse embryonic palatal mesenchyme (MEPM) cells from E13.5 wildtype and *Specc1l^ΔCCD2/ΔCCD2^* mutant embryos (Goering et al., 2021). Time-lapse imaging of wildtype and mutant MEPM cell cultures revealed that ΔCCD2 mutant cells exhibited a statistically significant deficit in their ability to form streams, an attribute of collective cell motility (**Figure S4**). This deficit in collective cell motility was consistent with our previous studies in SPECC1L deficient MEPM cells (Goering et al., 2021). At the cellular level, wildtype SPECC1L showed a diffuse punctate cellular expression pattern (**Figure 5A**), which closely mimicked F-actin expression pattern (**Figure 5C**). In contrast, SPECC1L-ΔCCD2 protein showed an abnormal perinuclear aggregation (**Figure 5E**). This abnormal perinuclear aggregation was even more vivid in migrating MEPM cells in a wound repair assay (**Figure S5**). While the mutant protein still colocalized with F-actin (**Figure 5H**), there appeared to be a drastic decrease in F-actin staining in the perinuclear region where the SPECC1L-ΔCCD2 protein had aggregated (**Figure 5E,G,H**). Superresolution STED imaging revealed a general association of both WT (**Figure 5K)** and mutant (**Figure 5O)** SPECC1L expression and actin filaments with reduced intensity. This was consistent with our previous finding of increased actin filaments upon SPECC1L deficiency (Hall et al., 2020; Wilson et al.). The microtubule expression pattern in cultured wildtype and mutant MEPM cells did not appear altered (**Figure 5B** vs. **5F; 5L** vs. **5P**).

**Figure 5.**
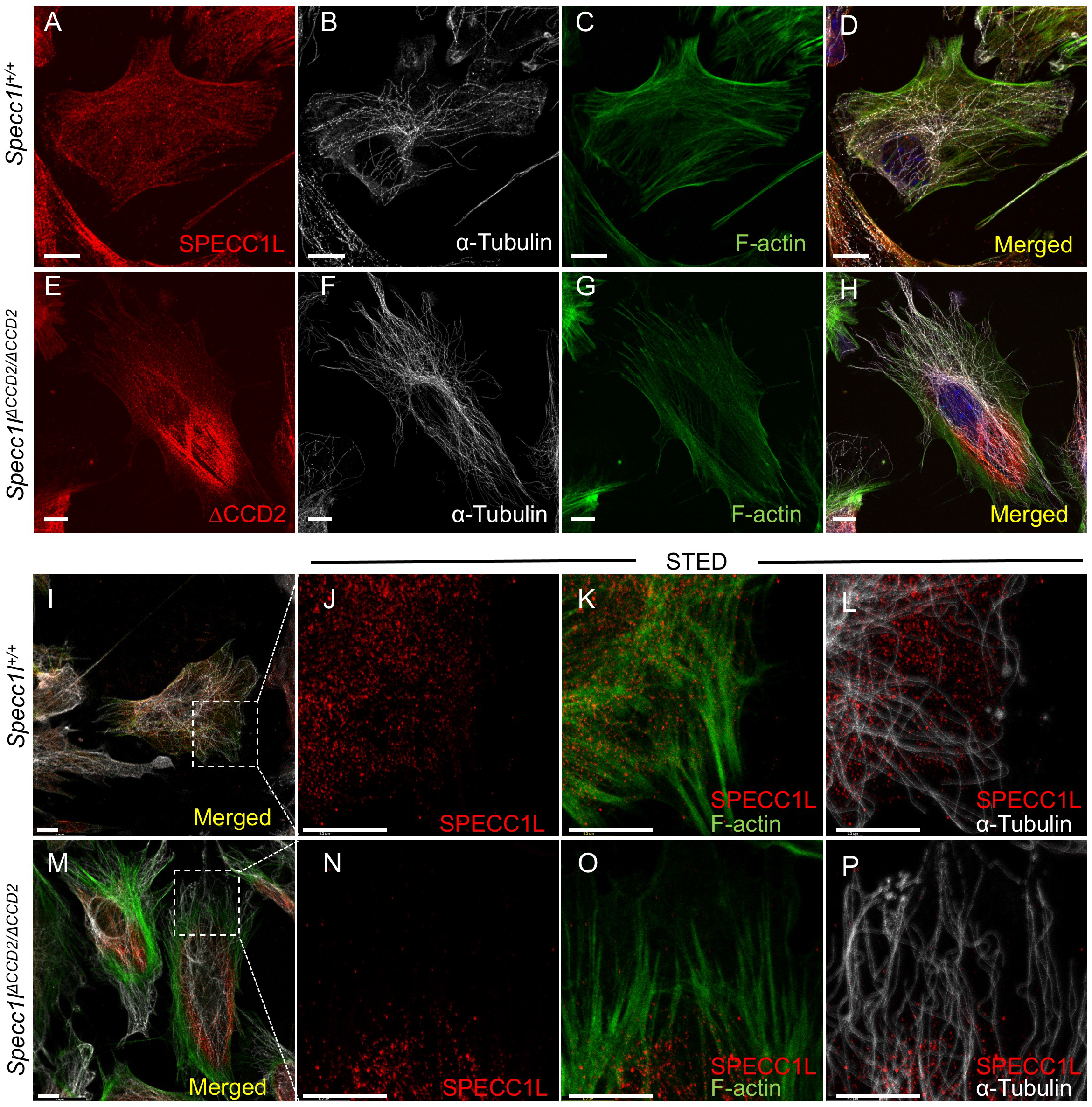
SPECC1L-ΔCCD2 shows abnormal perinuclear localization in primary mouse embryonic palatal mesenchyme cells. Primary mouse embryonic palatal mesenchyme (MEPM) cells were isolated from wildtype (A-D, I-L) and *Specc1l^ΔCCD2/ΔCCD2^* mutant (E-H, M-P) embryonic day (E) 13.5 embryos. MEPM cells were co-stained with antibodies against SPECC1L and a-Tubulin, and with phalloidin as indicated. Merged images with DAPI are also shown (D, H). SPECC1L shows a diffused subcellular expression pattern in wildtype MEPMs (A, D) that greatly overlaps with both microtubules (B, D) and actin filaments (F-actin; C, D). In contrast, ΔCCD2 protein in mutant MEPMs is clustered in a perinuclear region (E, H) with significantly reduced overlap with either microtubules (F, H) or F-actin (G, H). To better characterize the co-staining, superresolution STED images were taken (J-L, N-P) of the boxed regions shown in panels I (wildtype) and M (ΔCCD2 mutant). STED images show a strong overlap between SPECC1L and F-actin (K, O). Interestingly, regions of SPECC1L overlap show reduced F-actin staining. With microtubules, wildtype SPECC1L shows considerable overlap (L) compared to ΔCCD2 mutant (P). However, the main difference is an inability of mutant ΔCCD2 protein to distribute throughout the cytoplasm. Scalebars = 10 μm.

However, in contrast to WT MEPM cells (**Figure 5L**), there are clear regions of microtubules devoid of SPECC1L protein in mutant MEPM cells (**Figure 5P**). Thus, we propose that SPECC1L protein requires its CCD2 based association with microtubules for intracellular transport and diffusion. Furthermore, abnormal subcellular aggregation of SPECC1L-ΔCCD2 protein largely affected the actin cytoskeleton organization.

### Abnormal staining of non-muscle myosin II in *Specc1l^ΔCCD2/ΔCCD2^* MEPM cells

Non-muscle myosin II (NM-II) family consists of three types - A, B, and C - that are distinguished by three heavy chains encoded by *Myh9* (NM-IIA), *Myh10* (NM-IIB), and *Myh14* (NM-IIC), respectively (Ma & Adelstein, 2014b; Newell-Litwa, Horwitz, & Lamers, 2015). While NM-IIC is not expressed in the palate during development, both NM-IIA and NM-IIB have been implicated in palatogenesis (Kim et al., 2015) and the etiology of cleft palate (Birnbaum et al., 2009; Chiquet et al., 2009; Ma & Adelstein, 2014a). We looked at expression of both MYH9 and MYH10 in wildtype and *Specc1l^ΔCCD2/ΔCCD2^* mutant MEPM cells. We observed that both MYH9 (**Figure 6A,B**) and MYH10 (**Figure 6C,D**) expression pattern was abnormal in mutant MEPM cells. There was a particularly increased localization of both NM-IIA and NM-IIB to cell periphery. There was also a change in cell shape, with many mutant MEPM cells being larger and more circular (**Figure 6B,D**). We also extended our analysis to the palate mesenchyme (**Figure S6**) and found a similarly altered expression to the cell periphery of both MYH9 (**Figure S6A** vs. **S6F**) and MYH10 (**Figure S6K** vs. **S6P**). There also appeared to be reduced colocalization with microtubules of both NM-IIA (**Figure S6D** vs. **S6I, arrows**) and NM-IIB (**Figure S6N** vs. **S6S, arrow**).

**Figure 6.**
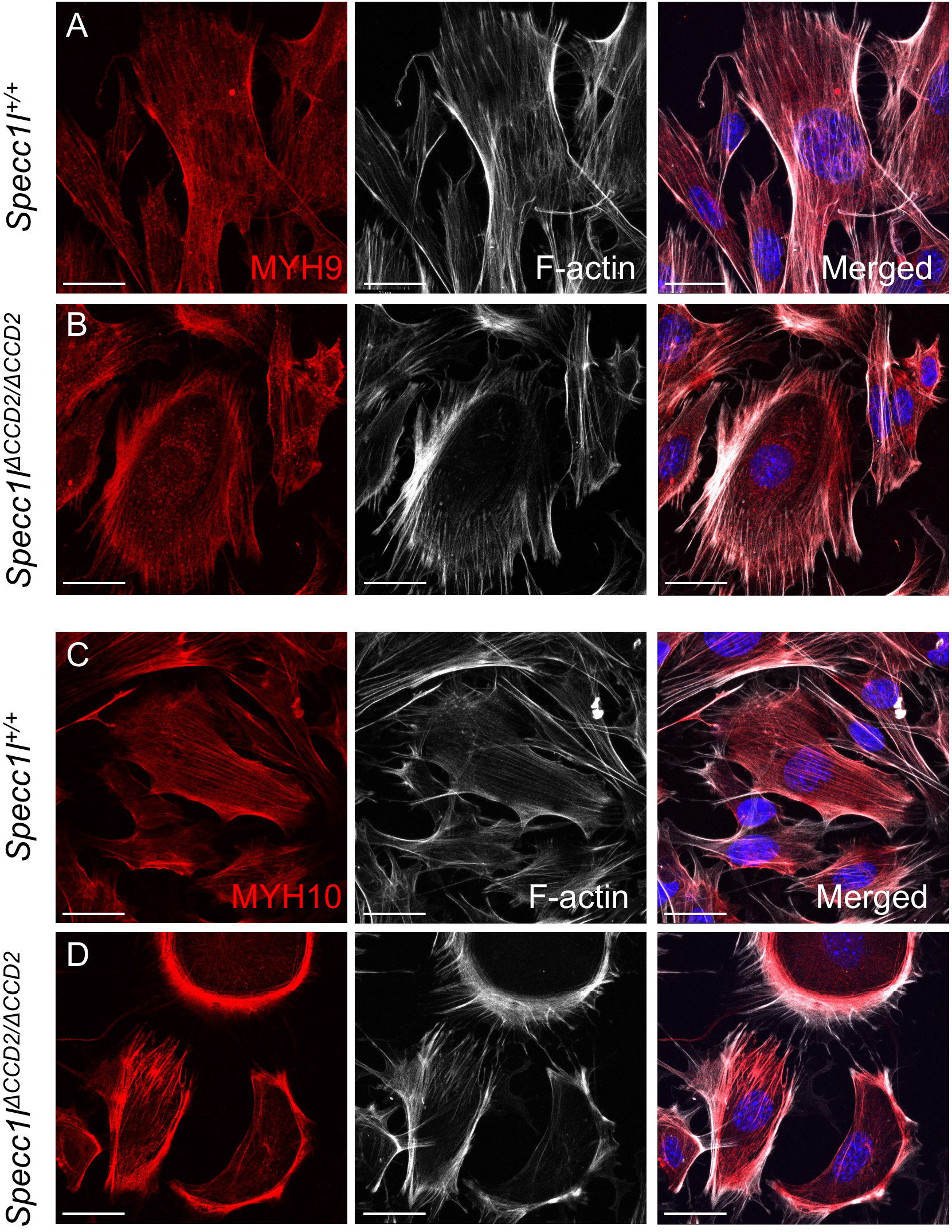
Abnormal non-muscle Myosin II expression in *Specc1l^ΔCCD2/ΔCCD2^* mutant primary mouse embryonic palatal mesenchyme cells. Wildtype (A, C) and *Specc1l^ΔCCD2/ΔCCD2^* mutant (B, D) primary mouse embryonic palatal mesenchyme (MEPM) cells were co-stained with antibodies against non-muscle myosin IIA (NM-IIA) component MYH9 (A, B) or with NM-IIB component MYH10 (C, D), and with phalloidin marking filamentous actin (F-actin). Merged images are shown with DAPI. Wildtype MYH9 (A) and MYH10 (C) show a uniform subcellular expression pattern that overlaps F-actin staining. In contrast, both MYH9 (B) and MYH10 (D) show an abnormal clustered expression at the cell periphery along with F-actin. Compared with wildtype MEPM cells (A, C), the mutant MEPM cells also show a more rounded appearance (B, D). Scalebars = 10 μm.

### NM-IIB coimmunoprecipitates with SPECC1L

Given the altered expression pattern of NM-IIA and NM-IIB in *Specc1l^ΔCCD2/ΔCCD2^* mutant MEPM cells and tissue, we wanted to know if wildtype or mutant SPECC1L associated with NM-II. We immunoprecipitated SPECC1L protein from wildtype, *Specc1l^ΔCCD2/ΔCCD2^* and *Specc1l^ΔC510/ΔC510^* embryonic tissue (**Figure 7**). Wildtype SPECC1L strongly coimmunoprecipitated NM-IIB (**Figure 7A**), as well as NM-IIA (not shown), showing an association *in vivo.* Interestingly, *Specc1l^ΔCCD2/ΔCCD2^* mutant protein also co-immunoprecipitated NM-IIB (**Figure 7B**), indicating that its association with SPECC1L was not via CCD2. Therefore, we tested lysate from *Specc1l^ΔC510/ΔC510^* mutants lacking the C-terminal CHD. The SPECC1L-ΔC510 protein also coimmunoprecipitated NM-IIB (**Figure 7C**), suggesting that there was a novel interaction domain for NM-IIB likely in the N-terminal region of SPECC1L.

**Figure 7.**
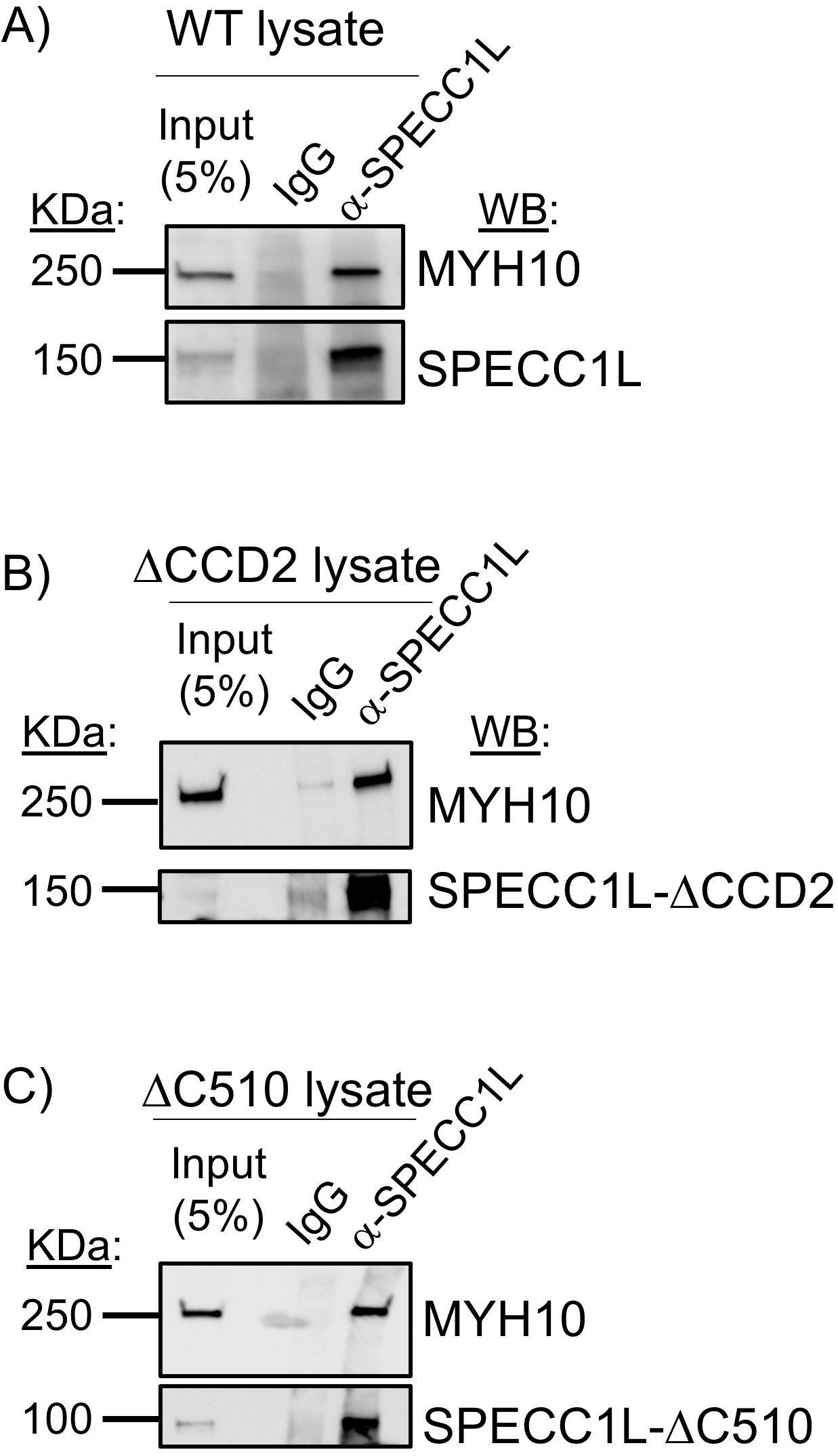
SPECC1L associates with NM-IIB component MYH10. Wildtype (A), *Specc1l^ΔCCD2/ΔCCD2^* mutant (B) and *Specc1l^ΔC510/ΔC510^* mutant (C) tissue lysate was used in coimmunoprecipitation assays using a-SPECC1L N-term antibody or IgG negative control. 5% input is shown as control. MYH10 was co-immunoprecipitated from WT, ΔCCD2 (lacking CCD2 domain) mutant and ΔC510 (lacking C-terminal CHD) mutant lysates. Thus, SPECC1L interaction with NM-IIB is not via CCD2 or CHD, but rather via a novel, yet unidentified N-terminal domain. WB = western blot.

## Discussion

Patients with autosomal dominant *SPECC1L* mutations show several structural birth defects including omphalocele and cleft palate (Bhoj et al., 2018). Yet in mice, we showed that a loss of the C-terminal region with CHD (Hall et al., 2020) or even a complete absence of SPECC1L protein (**Figure 1**) did not result in these defects at birth. We hypothesized that these autosomal dominant mutations that largely cluster in CCD2 are gain-of-function. We generated a mouse allelic series with in-frame deletions in CCD2 (**Figure 1**). Homozygous mutants for these in-frame ΔCCD2 deletions showed highly penetrant omphalocele and cleft palate, confirming our hypothesis that CCD2 perturbations are gain-of-function.

Furthermore, a significant proportion of ΔCCD2 heterozygous mutants also showed cleft palate, suggesting a dominant-negative mechanism. Interestingly, all ΔCCD2 heterozygous mutants with cleft palate were female. This is important since in humans, cleft palate only phenotype is observed more often in females (~1.41 times) than in males (Michalski et al., 2015; Rittler, Lopez-Camelo, & Castilla, 2004). We also observed an effect of background strain on the survival of ΔCCD2 heterozygotes, especially females. Together, these results indicate that our mouse *Specc1l^ΔCCD2^* alleles aptly model the human condition.

Homozygous in-frame ΔCCD2 mutants also showed highly penetrant exencephaly and coloboma. While cleft palate and omphalocele has been frequently observed in known patients with *SPECC1L* mutations, exencephaly and coloboma are not. Coloboma was only observed in a small subset of homozygous mutants, and not in heterozygous mutants, thus suggesting that optic fissure closure may not be sensitive to one copy of the CCD2 mutation. In contrast the main reason for absence of exencephaly in the patients identified to-date is likely selection bias. The current patient cohort was initially identified based on *SPECC1L* association with craniofacial anomalies (Kruszka et al., 2015; Saadi et al., 2011) and then with omphalocele (Bhoj et al., 2015). Fetuses with exencephaly fail to survive postnatally and are not currently associated with *SPECC1L* mutations. Since we do observe occurrence of exencephaly in our heterozygous mouse mutants, we expect that *SPECC1L* mutations will eventually be identified in genetic studies specifically focused on exencephaly or perhaps other neural tube defects.

SPECC1L appears to affect the efficiency of several embryonic tissue movement and fusion events. Several pieces of evidence for this efficiency argument come from our study of palatogenesis in various *Specc1l* alleles. First is the incomplete penetrance of the various fusion defects in ΔCCD2 mutants. Another is that in SPECC1L truncation (Hall et al., 2020) and null (**Figure 2**) alleles there is no cleft palate at birth, indicating that SPECC1L is not required for the process, but there is a demonstrable delay in palatal shelf elevation (Hall et al., 2020). We hypothesize that this elevation delay is more severe in in-frame ΔCCD2 mutants, which results in cleft palate at birth. A good case in point for this hypothesis is the absence of cleft palate in ΔCCD2 mutant embryos with exencephaly where the oral cavity is narrowed due to the loss of the cranium. Even with reduced efficiency in elevation, palatal shelves have a shorter distance to cover and are thus able to fuse. An important corollary to this argument is that even subtle variants in *SPECC1L* may affect this efficiency enough to function as genetic modifiers. This is consistent with our identification of *SPECC1L* variants in patients with non-syndromic cleft lip/palate with subtle functional consequences (Hall et al., 2020) compared to variants identified in syndromic cases (Kruszka et al., 2015; Saadi et al., 2011). We propose that the same would be true in other fusion defects of the neural tube, ventral body wall and eye.

The basis for this reduced efficiency in the SPECC1L-deficient palate is likely poor vertical to horizontal remodeling during elevation (Goering et al., 2021). Several cellular mechanisms for this mesenchymal remodeling have been proposed (Bush & Jiang, 2012; Gritli-Linde, 2008; Lan, Xu, & Jiang, 2015; Li, Lan, Krumlauf, & Jiang, 2017). Cell proliferation is required, but not sufficient (Jin, Li, Higashi, Darling, & Ding, 2008; Kouskoura et al., 2013; Lan, Zhang, Liu, Xu, & Jiang, 2016), however, migratory properties, potentially guided by WNT5A and FGF10 chemotactic gradients (He et al., 2008) may be needed for elevation to occur. We posited that, at the cellular level, SPECC1L deficiency leads to poor cell alignment and motility (Goering et al., 2021), two key elements of coordinated movement that can affect this remodeling. At the molecular level, the ability of SPECC1 L to associate with both microtubules and actin via CCD2 and CHD, respectively, underlies its function. We had hypothesized that CCD2 based association with microtubules was critical, as most human mutations clustered in this domain, however, the significance of the microtubule association was not evident. Previous *in vitro* assays suggested an ability to promote acetylation or stability of a subset of microtubules (Hall et al., 2020; Saadi et al., 2011). Our data now show that while there is some reduction or mislocalization of microtubules in ΔCCD2 mutants, the main effect of CCD2 loss is an inability of SPECC1L to evenly distribute within the cell cytoplasm. Thus, SPECC1L is likely using microtubule-based trafficking to move throughout the cell. Furthermore, the subcellular regions, where the mutant SPECC1L-ΔCCD2 protein aggregates, are conspicuously devoid of strongly staining actin filaments. This observation is consistent with a role for SPECC1L in actin depolymerization, and with our previous reports of increased F-actin staining in SPECC1L-deficient cells (Hall et al., 2020; Wilson et al., 2016).

Given our findings with actin cytoskeletal changes, we looked at NM-IIA and NM-IIB. NM-IIA (*MYH9*) has been implicated in association studies with nonsyndromic CL/P (Birnbaum et al., 2009; Chiquet et al., 2009; Ma & Adelstein, 2014a) and in palatogenesis (Kim et al., 2015). Interestingly, similarly to *Specc1l,* NM-IIB (*Myh10*) null allele did not show cleft palate or omphalocele (Ma & Adelstein, 2014b), while an *Myh10* allele with a point mutation showed cleft palate, omphalocele and other structural birth defects (Ma & Adelstein, 2014a). We observed an abnormal NM-IIA and IIB staining pattern towards the periphery of mutant MEPM cells, likely associated with branched actin filaments. We also observed a general rounding of mutant MEPM cells, which is consistent with a decrease in cytoplasmic association of NM-II with central long anti-parallel actin filaments. Most importantly, we show that SPECC1L physically associates with NM-II, independent of either the CCD2 (microtubule) or CHD (actin) domains. This result further establishes SPECC1L as a cytoskeletal scaffolding protein.

By organizing cellular actin and myosin, SPECC1L may participate in the global alignment of actomyosin based forces that play a critical role in time-sensitive embryonic tissue movement and fusion events.

## Acknowledgements

This project was supported in part by the National Institutes of Health grants DE026172 (I.S.), GM102801 (A.C.), and F31DE027284 (E.H.). I.S. was also supported in part by the Center of Biomedical Research Excellence (COBRE) grant (National Institute of General Medical Sciences P20 GM104936), Kansas IDeA Network for Biomedical Research Excellence grant (National Institute of General Medical Sciences P20 GM103418), and Kansas Intellectual and Developmental Disabilities Research Center (KIDDRC) grant (U54 Eunice Kennedy Shriver National Institute of Child Health and Human Development, HD090216). The Confocal Imaging Facility, the Integrated Imaging Core, and the Transgenic and Gene Targeting Institutional Facility at the University of Kansas Medical Center are supported, in part, by NIH/NIGMS COBRE grant P30GM122731 and by NIH/NICHD KIDDRC grant U54HD090216. The Leica STED microscope is supported by NIH 1S10OD023625.

## Author contributions

JPG, LWW, MS, AC, and IS conceived and designed the experiments. JPG, LWW, MS, MM, EGH, DGI, BJ, ZU and MKR performed the experiments. DGI and AC performed the time-lapse imaging analysis. JPG, MS, AC and IS wrote the paper. LWW, MM, BJ and MKR edited the manuscript. All authors reviewed the manuscript.

## Conflict of Interest

The authors do not have any competing financial interests pertaining to the studies presented here.

## Notes

### Competing Interest Statement

The authors have declared no competing interest.

