## Supplemental data for "In-frame deletion of SPECC1L microtubule binding domain results in embryonic tissue movement and fusion defects"

Supplemental Table 1

**Table S1: Penetrance of exencephaly, cleft palate and omphalocele phenotypes in *Specc1l* alleles.**

| Mouseline | Genotype | n | Exencephaly | Cleft Palate | Omphalocele |
|---|---|---|---|---|---|
| Δ234 | *Specc1l*$^{+/+}$ | 20 | 0.0 | 0.0 | 0.0 |
| | *Specc1l*$^{\Delta 234/+}$ | 31 | 6.5 | 9.7 | 3.2 |
| | *Specc1l*$^{\Delta 234/\Delta 234}$ | 14 | 57.1 | 35.7 | 71.4 |
| Δ576 | *Specc1l*$^{+/+}$ | 8 | 0.0 | 0.0 | 0.0 |
| | *Specc1l*$^{\Delta 576/+}$ | 18 | 5.6 | 0.0 | 22.2 |
| | *Specc1l*$^{\Delta 576/\Delta 576}$ | 14 | 28.6 | 50.0 | 50.0 |
| Δ411 | *Specc1l*$^{+/+}$ | 5 | 0.0 | 0.0 | 0.0 |
| | *Specc1l*$^{\Delta 411/+}$ | 18 | 0.0 | 5.6 | 16.7 |
| | *Specc1l*$^{\Delta 411/\Delta 411}$ | 6 | 0.0 | 16.7 | 83.3 |
| ΔEx4 | *Specc1l*$^{+/+}$ | 15 | 0.0 | 0.0 | 0.0 |
| | *Specc1l*$^{\Delta Ex4/+}$ | 33 | 0.0 | 0.0 | 0.0 |
| | *Specc1l*$^{\Delta Ex4/\Delta Ex4}$ | 16 | 0.0 | 0.0 | 0.0 |

### Supplemental Figure 1

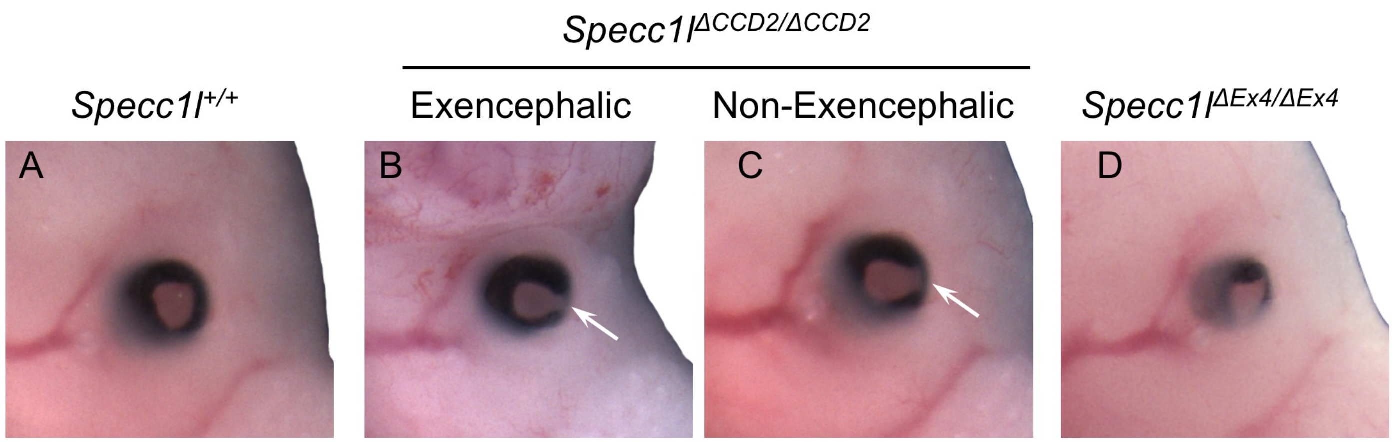


**Figure S1: Coloboma in *Specc1l*$^{\Delta CCD2/\Delta CCD2}$ mutants.** *Specc1l*$^{\Delta CCD2/\Delta CCD2}$ mutants either with exencephaly (B) or without exencephaly (C) frequently show incomplete closure of the optic or choroid fissure, resulting in coloboma (arrows). This defect is not observed in wildtype (A) or *Specc1l*$^{\Delta Ex4/\Delta Ex4}$ mutants (D).

Supplemental Figure 2

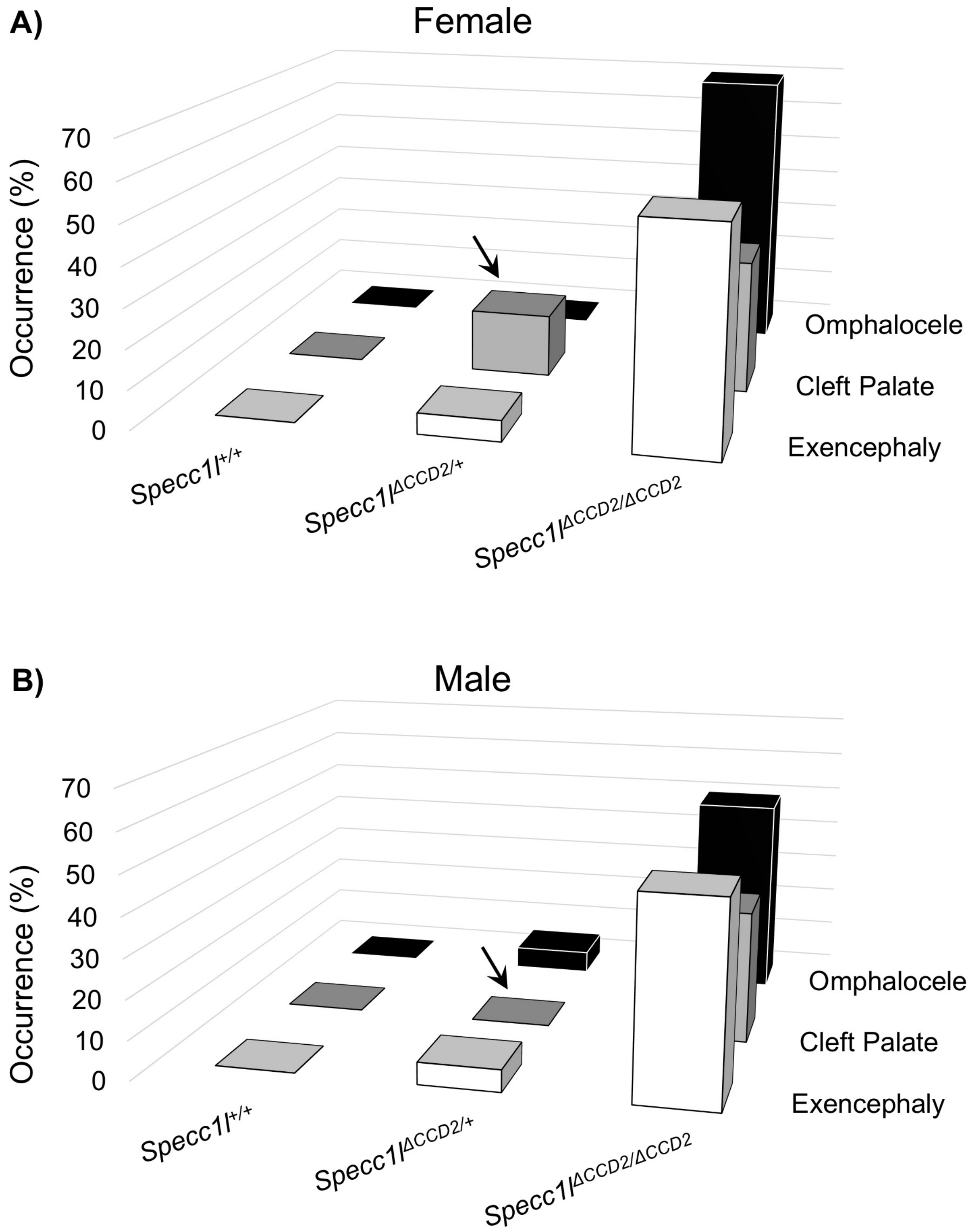


**Figure S2: Increased occurrence of cleft palate in female *$Specc1l^{\Delta CCD2/+}$* heterozygotes.** The penetrance of omphalocele, cleft palate and exencephaly shown in Figure 1 and Supplemental Table 1 was assessed separately in female **(A)** and male **(B)** embryos. The three phenotypes occur similarly in *$Specc1l^{\Delta CCD2/\Delta CCD2}$* homozygous mutants. However, among *$Specc1l^{\Delta CCD2/+}$* heterozygotes, cleft palate was observed almost exclusively in female embryos **(arrows)**. There is also an increased occurrence of omphalocele in male heterozygotes. Exencephaly occurs similarly again in female and male heterozygotes.

Supplemental Figure 3

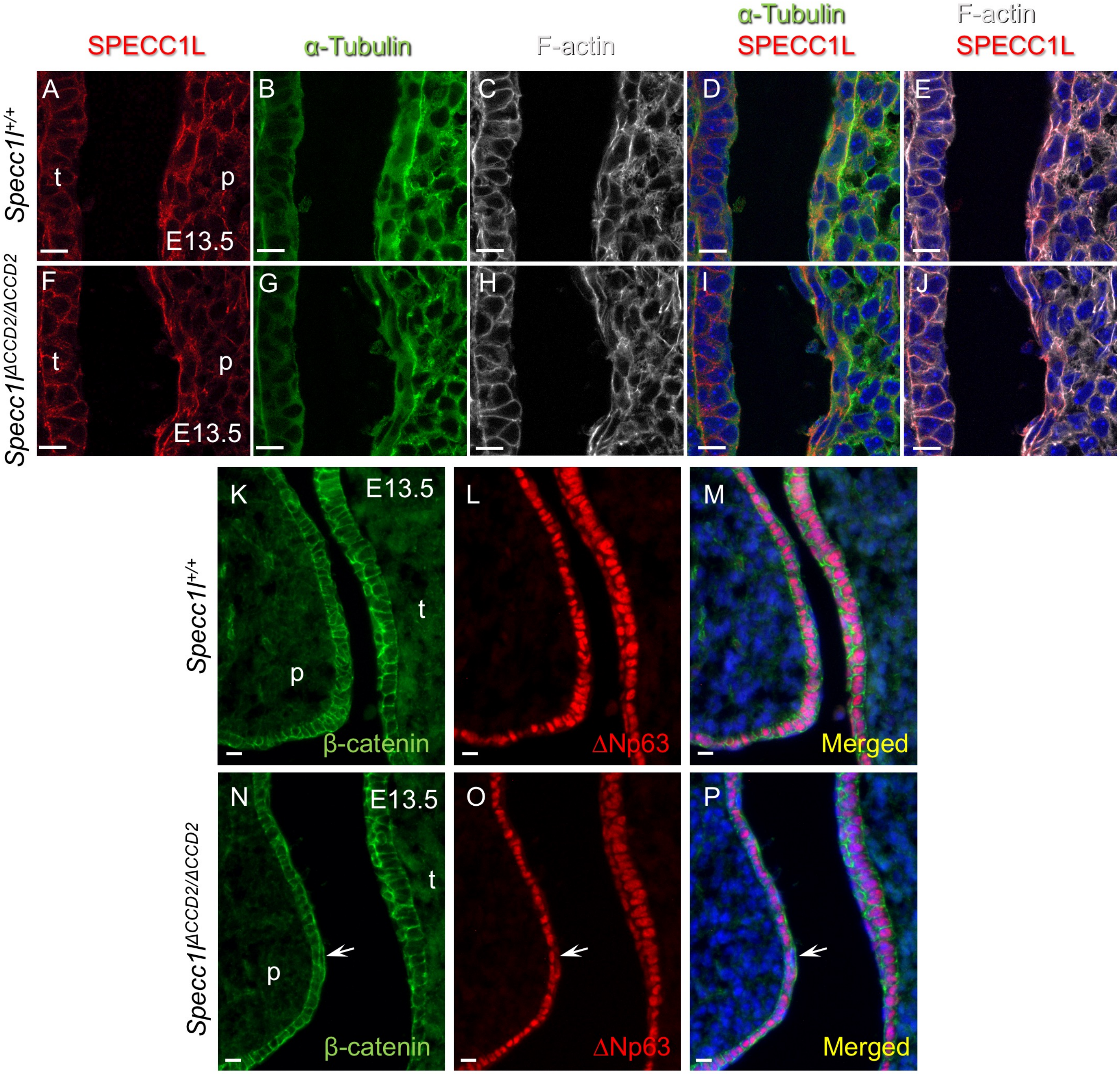


**Figure S3: Cellular expression of SPECC1L-ΔCCD2 and β-catenin in E13.5 palatal shelf epithelium. A-J)** Wildtype *Specc1l*$^{+/+}$ (A-E) and *Specc1l*$^{\Delta CCD2/\Delta CCD2}$ mutant (F-J) embryonic day (E) 13.5 cryosections were co-stained with antibodies against SPECC1L (A, F) and α-Tubulin (B, G), and with phalloidin (C, H). Merged images of wildtype (D, E) and mutant (I, J) SPECC1L with microtubules (D, I) or actin filaments (F-actin; E, J) are shown with DAPI. The abnormal localization of mutant protein to the cell periphery (A vs. F) is less obvious in the palatal shelf epithelium than in the mesenchyme (Figure 4). However, compared to wildtype (D), there is still a reduced overlap between mutant ΔCCD2 protein and microtubules (I). **K-P)** Widltype (K-M) and *Specc1l*$^{\Delta CCD2/\Delta CCD2}$ mutant (N-P) E13.5 sections were also co-stained with antibodies against β-catenin (K, N), marking cell-cell adhesions, and ΔNP63 (L, O), marking the basal epithelium. Merged images are shown with DAPI (M, P). Flat ΔNP63-negative cells on the epithelium surface are periderm cells that do not express adhesion molecules like β-catenin on their apical surface (M). In the *Specc1l*$^{\Delta CCD2/\Delta CCD2}$ mutant, only rare occurrences were found of abnormal apical expression of β-catenin on the apical surface of ΔNP63-negative periderm cells (P, arrow). Scalebars = 10μm; p = palatal shelf; t = tongue.

### Supplemental Figure 4

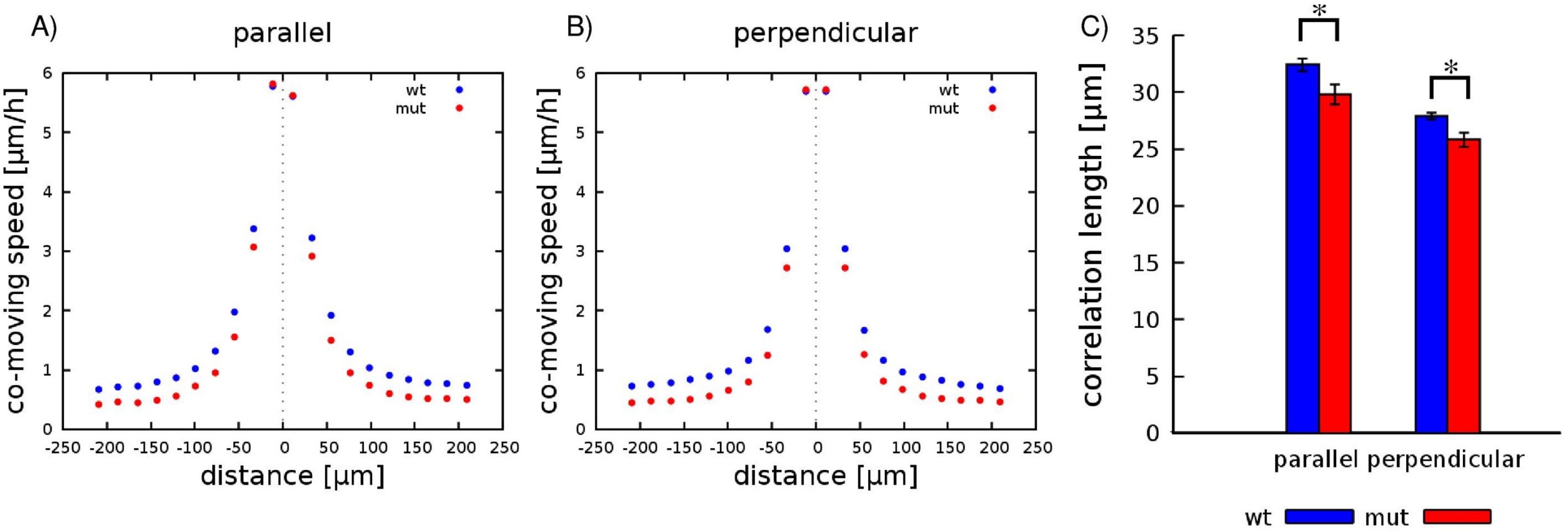


**Figure S4: Primary mouse embryonic palatal mesenchyme cells from *Specc1l*$^{\Delta CCD2/\Delta CCD2}$ mutant showed poor directional cell stream formation.** Primary MEPM cells from wildtype (wt) and *Specc1l*$^{\Delta CCD2/\Delta CCD2}$ mutant (mut) embryos exhibited directional alignment and collective motility in high cell density 2D motility assays. MEPM cells tended to move in the same direction locally, indicated by positive average co-moving speed values -- both along the front-rear (parallel, A) and left-right (perpendicular, B) axes. The co-movement of *Specc1l*$^{\Delta CCD2/\Delta CCD2}$ mutant cells, however, extended to a shorter distance than that of WT cells. C) Average correlation lengths, both parallel with and perpendicular to the direction of the local prevailing direction of cell motion, were established by fitting an exponential function on the profiles shown in panels (A, B). Data were averaged from 4 independent fields, both for WT (red) and *Specc1l*$^{\Delta CCD2/\Delta CCD2}$ mutant (blue) cells. Asterisks denote statistical significance (p=0.02 and p=0.008 for the parallel and perpendicular directions, respectively).

### Supplemental Figure 5

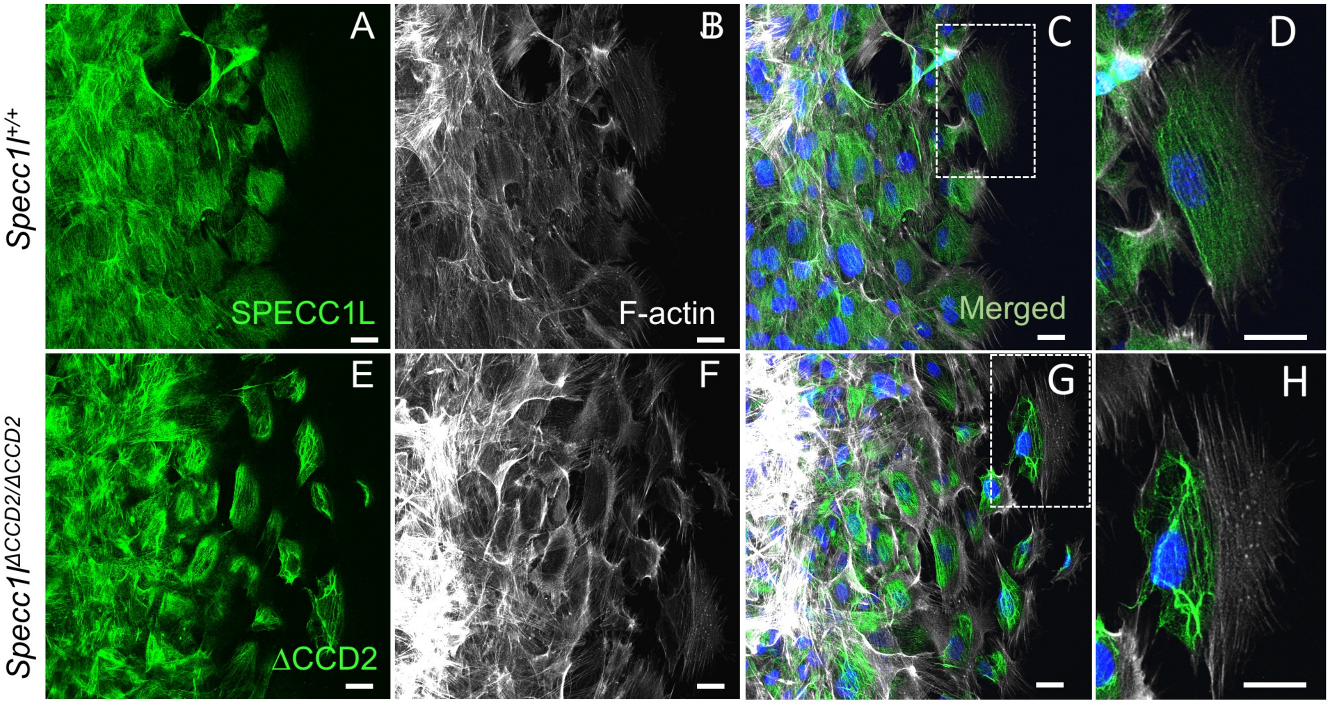


**Figure S5: SPECC1L-ΔCCD2 shows abnormal perinuclear localization in migrating primary mouse embryonic palatal mesenchyme cells.** Primary mouse embryonic palatal mesenchyme (MEPM) cells were isolated from wildtype (A-D) and *Specc1l*$^{\Delta CCD2/\Delta CCD2}$ mutant (E-H) embryonic day (E) 13.5 embryos. For immunostaining, MEPM cells were captured in the process of migration in a wound-repair assay. In panels A-H, migrating MEPMs were co-stained with α-SPECC1L (A, E) and phalloidin (B, F). Merged images with DAPI (C, G) and a magnification of the boxed regions are also shown (D, H). Migrating wildtype MEPMs show diffused SPECC1L expression (A, C, D) that closely matches that of F-actin (B, C, D). In contrast, ΔCCD2 protein in mutant MEPMs is expressed in abnormal bundles in a perinuclear region confined to the posterior end of the migrating cell (E, G, H). Scalebars = 25 μm.

Supplemental Figure 6

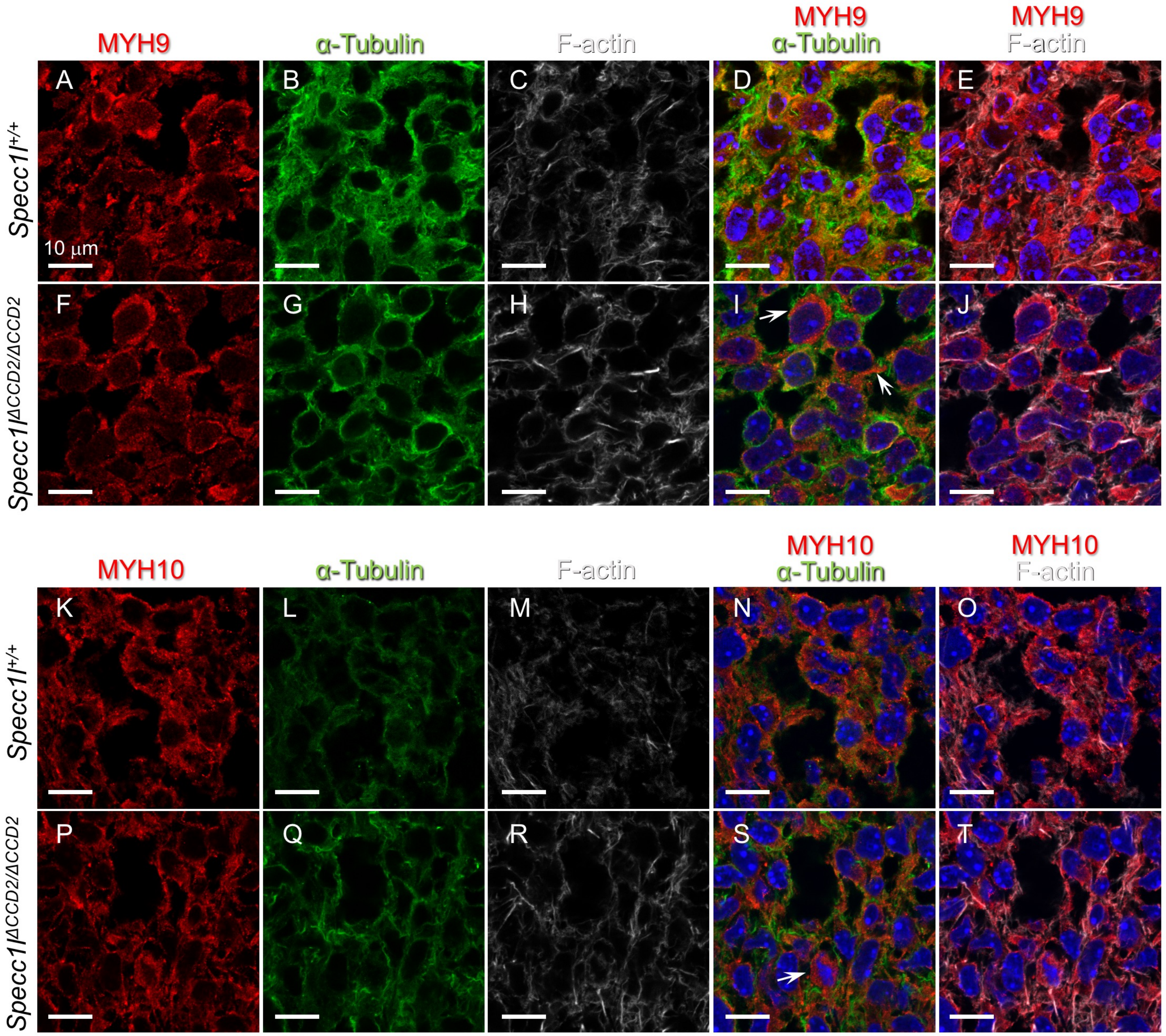


**Figure S6: Abnormal non-muscle Myosin II expression in *Specc1l$^{\Delta CCD2/\Delta CCD2}$* mutant palatal shelf mesenchyme.** Wildtype (A-E, K-O) and *Specc1l$^{\Delta CCD2/\Delta CCD2}$* mutant (F-J, P-T) cryosections through the palatal shelf mesenchyme were co-stained with antibodies against non-muscle myosin IIA (NM-IIA) component MYH9 (A, F) or with NM-IIB component MYH10 (K, P), and α-Tubulin (B, G, L, Q), and with phalloidin marking filamentous actin (F-actin; C, H, M, R). Merged images of MYH9 (D, E, I, J) or MYH10 (N, O, S, T) with microtubules (D, I, N, S) or F-actin (E, J, O, T) are shown with DAPI. Wildtype MYH9 (D) and MYH10 (N) show a subcellular expression pattern that overlaps well with microtubule staining. In contrast, in *Specc1l$^{\Delta CCD2/\Delta CCD2}$* mutant tissue, both MYH9 (I) and MYH10 (J) show a reduced overlap with microtubule staining (arrows). Scalebars = 10 μm.
